# A mosaic of climate vulnerability: local warming rates meet intraspecific divergence in heat tolerance

**DOI:** 10.64898/2026.03.20.713229

**Authors:** Wissam A. Jawad, Ana L. Salgado, Brian S. Cheng, Sarah A. Gignoux-Wolfsohn, Cynthia Hays, Martha M. Muñoz, Matthew C. Sasaki, Morgan W. Kelly

**Affiliations:** Department of Biological Sciences, Louisiana State University, Baton Rouge, LA 70808; Department of Environmental Conservation, University of Massachusetts, Amherst, MA, 01003; Department of Biological Sciences, University of Massachusetts, Lowell, MA, 01854; Department of Biological Science, Keene State College, Keene, NH, 03435; Department of Ecology and Evolutionary Biology, Yale University, New Haven, CT, 06520

**Author notes:** **Corresponding author:** Wissam Jawad. Joint first authors on this manuscript (Wissam A. Jawad, Ana L. Salgado). **Statement of authorship:** All authors conceived the idea and designed the general study. A.L.S and W.A.J developed the methods for processing and analyzing data with assistance from B.S.C, M.C.S, and M.W.K. W.A.J wrote the first draft of the manuscript. All authors contributed to revising and editing the manuscript. **Data accessibility** All data and code can be accessed, upon publication, via Zenodo.

**Keywords:** climate warming, heat tolerance, local adaptation, thermal safety margin, warming tolerance, evolutionary rescue, CT max, upper thermal limit

## Abstract

Climate warming is increasing mismatches between thermal phenotypes and habitat temperatures, driving range shifts and population extirpations. While within-species variation in heat tolerance and local warming rates can predict responses to climate warming, how these factors shape differences in vulnerability among taxa and ecosystems is uncertain. Here we combine climate and thermal trait data from 69 species across four ecosystem types to examine the effects of incorporating intraspecific variation in heat tolerance and local warming rates on projected vulnerability to climate warming. Because vulnerability to warming depends on existing phenotypic variation in thermal performance and relative rates of habitat warming, we develop a new metric that integrates localized rates of warming with spatial variation in thermal tolerances, termed the ‘minimum trait velocity’. Incorporating intraspecific variation in heat tolerance lowered estimates of warming tolerance (a measure of vulnerability) across most ecosystem types, with the strongest negative impact on marine taxa. Although intraspecific variation in heat tolerance could facilitate adaptation to climate change, our results suggest such variation is generally less than the projected near future warming. This suggests that opportunities for evolutionary rescue via gene flow between locally adapted populations are limited, adding to mounting concern as the climate warms.

## Introduction

Climate warming is causing variable and often unpredictable biological responses, including range expansions and local extirpations, shaped in part by the relationship between each species’ thermal niche and changing habitat temperatures (Bozinovik & Pörtner 2015, Wernberg et al. 2016, da Silva & Diamond 2024). Data on the match between species’ phenotypic responses to heat stress and their current and future habitat temperatures can provide invaluable insight for forecasting vulnerability to warming (Evans et al. 2015). However, most thermal niches are only characterized at the species level, which may under or overestimate species vulnerability relative to estimates based on more fine-scale data. Many species’ geographic ranges span wide environmental gradients, with populations physiologically adapted to local climatic conditions (Kuo & Sanford 2009, Kelly et al. 2011, Eliason et al. 2011, Dongmo et al. 2021). Across these climactically heterogeneous ranges, defaunation or the loss of populations will occur before extinction (Dirzo et al. 2014). Intraspecific analyses are needed to understand the process of defaunation because the effects of changes in climate are likely to be population-specific and nuanced by the relationship between each population’s current environmental tolerances and the rate of climate change at that location.

Recent studies have demonstrated the important but often ignored role of fine-scale features, such as intraspecific variation in thermal tolerance and local climate conditions, in shaping predictions of species vulnerability across broader taxonomic and regional scales (Angert et al. 2011, Bennet et al. 2019, Baker et al. 2025). Crucially, such considerations can fundamentally reconfigure predictions of vulnerability under climate warming. For example, projected areas of thermally suitable habitats can shrink when predictions include local adaptation within the species, as opposed to treating the species as a single entity with a single phenotypic response to warming (Valladers et al. 2014). In some cases, species previously predicted to experience greater habitat suitability under climate warming would experience more than 50% loss of suitable habitat when local adaptation was incorporated (Cacciapaglia & van Woesik 2018). Authors have also demonstrated the importance of including local variation in rates of warming in predicting when habitats will become thermally unsuitable for a particular species (da Silva & Diamond 2024). Local climate can be more indicative of broad-scale species’ responses than regional level projections of environmental change (Bay et al. 2019, da Silva & Diamond 2024). It is thus crucial that work aimed at predicting species responses to climate warming across ecosystem types (i.e., marine and terrestrial taxa) and regions incorporate population-level phenotypic variation as well as local change in habitat temperatures.

Warming tolerance, or the difference between maximum habitat temperature and an organism’s upper thermal limit, quantifies species- and population-level vulnerability to habitat warming (Deutch et al. 2008, Collin et al. 2018, Villeneuve et al. 2021) that can be used as a standardized metric for comparisons between taxa and ecosystem types (Deutch et al. 2008, Pinsky et al. 2019). Differences in species warming tolerance within a region can be a predictor of demographic responses to warming, with higher thermal tolerance corresponding to greater abundances in warming habitats (Birkett et al. 2018, Roeder et al. 2021). Although there has been considerable progress in predicting the effects of mismatches in phenotype and environment for specific taxa and habitats, data on variation in the magnitude of intraspecific variation in thermal tolerance is often missing when comparing vulnerability across broader scales, hampering our ability to predict how vulnerability differs across ecosystems.

The distribution of thermal tolerance phenotypes across populations also has important consequences for the probability of evolutionary rescue via gene flow. Warm-habitat populations often evolve higher thermal tolerances, providing a potential reservoir of adaptive genetic variation that could be shared between populations with lower thermal tolerance (Sanford and Kelly 2011, Somero 2010, Angert et al. 2011). Previous work has demonstrated that evolutionary rescue through gene flow can save populations experiencing changing environmental conditions (Bell 2013, Fitzpatrick et al. 2016). Hybridization with warm-adapted populations can improve survival under warming in cool-adapted populations (Griffiths et al. 2021, Lewis et al. 2024, Brauer et al. 2023), suggesting gene flow could rescue less tolerant populations from habitat warming. Identifying the amount and distribution of phenotypic variation in thermal traits among populations can therefore aid in forecasting the contexts in which evolutionary rescue through gene flow would be most likely. The distance and degree of gene flow between populations with divergent thermal phenotypes will be especially crucial for species and populations experiencing fast rates of habitat warming, where maximum habitat temperatures could surpass thermal limits (Fig. 1). It is thus essential to understand how the distribution of thermal traits across space relates to the rate of change in local climate through time and how this relationship varies among species and ecosystems.

**Figure 1.**
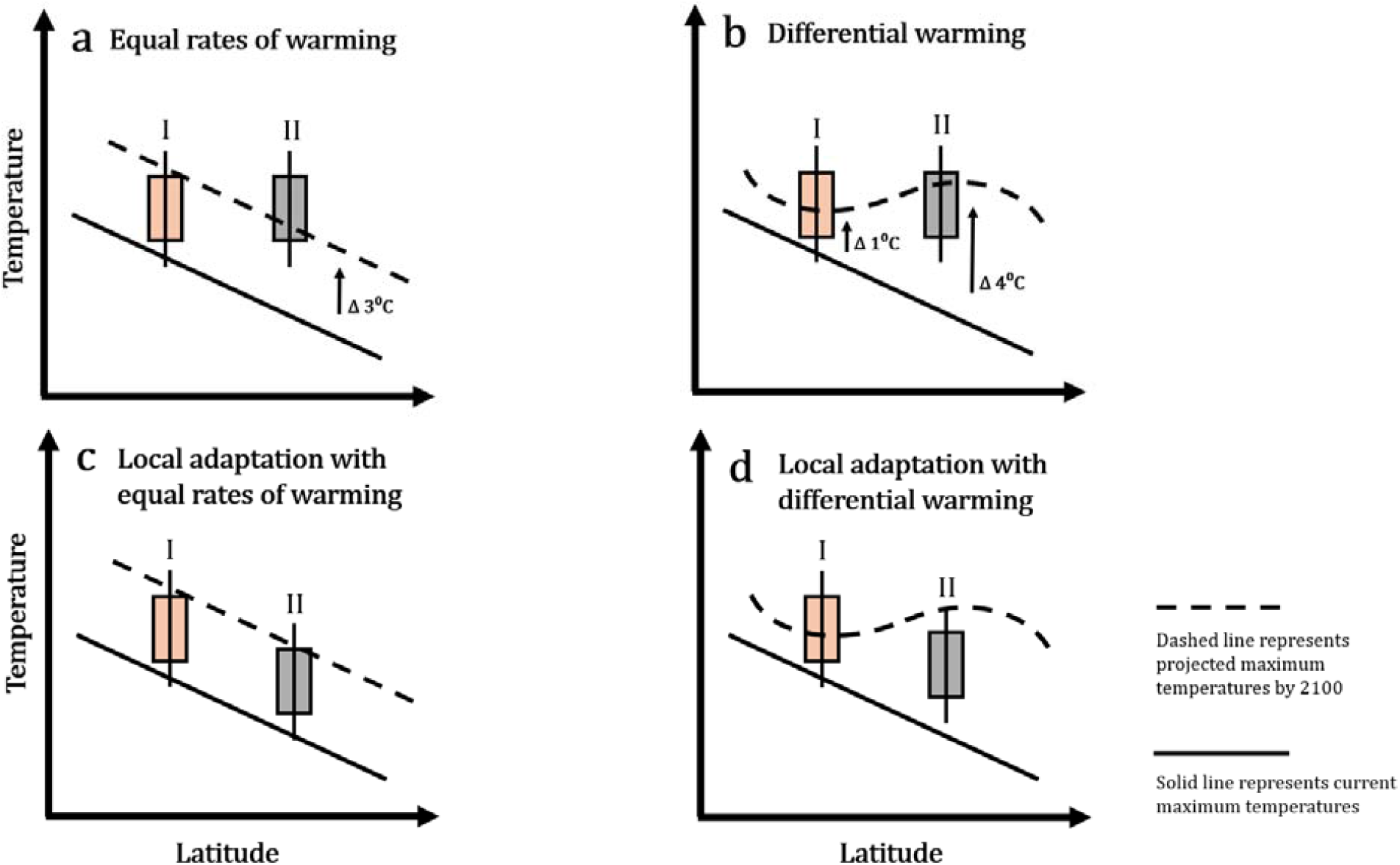
Implications for incorporating local adaptation and population-level rates of habitat warming for forecasting population vulnerability to warming (adapted from Sanford and Kelly 2011). Box plots represent the distribution of thermal tolerance phenotypes within two hypothetical populations (I and II) along a latitudinal temperature gradient. Solid black line represents the current maximum temperatures along the gradient, and the dashed line represents future maximum temperatures. Populations have the potential for persistence if some amount of the box plot distribution lies above the dotted line, while box plots completely below the dotted line indicate projected population extirpation. In (*a*), using a standard rate of habitat warming across two populations and without local adaptation, population I is projected to be more vulnerable to habitat warming than population II. In (*b*), using local rates of habitat warming for each population, both populations appear to persist, but now population II is most vulnerable. In (*c*), incorporating local adaptation but using a standard rate of habitat warming, both populations are highly vulnerability to extinction. In (*d*) by incorporating local rates of habitat warming with local adaptation, population I would persist, but population II would not. Note that these projections will also depend on the range of phenotypes present within each population, which could differ within and between taxa.

Comparisons of intraspecific variation in thermal traits among ecosystems have demonstrated that marine ectotherms harbor greater divergence across shorter spatial scales than do terrestrial ectotherms (Sasaki et al. 2022). Such a finding is surprising because the potential for local adaptation to climate gradients was thought to be lower in the sea due to high connectivity for species with planktonic larvae (but see Kelly & Sanford 2011). However, high connectivity in the sea appears to be outweighed by lower potential for buffering against selection via behavioral thermoregulation. In terrestrial systems, mobile ectotherms can minimize thermal stress through behavioral thermoregulation, effectively buffering against selection for higher thermal tolerance, a phenomenon termed the Bogert effect (Bogert 1949, Muñoz 2022). Marine and freshwater species, however, have limited capacity to behaviorally thermoregulate because there is less fine scale variation in temperature for organisms to seek thermal refugia. Thus, aquatic ectotherms are more directly exposed to selection for higher heat tolerance imposed by environmental temperatures than terrestrial species, resulting in greater local adaptation to temperature over shorter spatial scales and a potentially higher capacity for evolutionary rescue through gene flow compared to terrestrial ectotherms. However, whether adaptation can keep pace with climate change will also depend on the relative rates of warming between marine and terrestrial populations, making it essential to integrate local rates of warming with intraspecific thermal trait data.

Here we use a data synthesis approach to integrate two types of fine scale data to estimate climate vulnerability: adaptive differentiation in upper thermal limits within species and local projections of habitat warming. We use these data to evaluate the current and future population-level warming tolerances across marine, terrestrial, freshwater and intertidal ecosystems, and test whether projections of vulnerability are substantially altered by incorporation of population-level trait and climate data. Specifically, we test the hypothesis that incorporating local adaption should have a larger effect on estimates of warming tolerance in marine taxa (marine and intertidal ecosystems) compared to terrestrial taxa, based on the observation that marine taxa harbor more divergence in heat tolerance compared to terrestrial taxa (Sasaki et al. 2022). In addition, we compare projected rates of habitat warming at local scales to the spatial scale of intraspecific variation in thermal limits to determine the speed that migration of locally adapted phenotypes would have to occur for populations with lower thermal tolerance to be rescued by gene flow from populations with higher thermal tolerance, a metric we term ‘Minimum Trait Velocity’ (see Text Box 1 for an example). We test the hypothesis that marine taxa will have lower estimated minimum trait velocities, due to having greater divergence for heat tolerance over shorter distances compared to terrestrial taxa, suggesting higher potential for evolutionary rescue through gene flow. The potential for evolutionary rescue through gene flow between thermally divergent populations is contingent on is the existence of sufficient divergence in heat tolerance relative to the rate of habitat warming between populations. In other words, if the projected increase in habitat temperatures at a populations site is greater than the difference in heat tolerance between that population and surrounding populations, hybridization between populations would have minimal capacity to buffer the more vulnerable population against increasing habitat temperatures. Thus, we also evaluate the proportion of pair-wise population comparisons across all four ecosystems that have a difference in heat tolerance that’s greater than the projected increase in habitat temperature in the near future (25 years from current). Our methodology and results offer a novel advance in integrating intraspecific variation in thermal tolerance and population-level rates of warming for predicting climate vulnerability of populations and taxa across ecosystems.

### Text Box 1 Case study for estimating minimum trait velocity between populations of an intertidal copepod

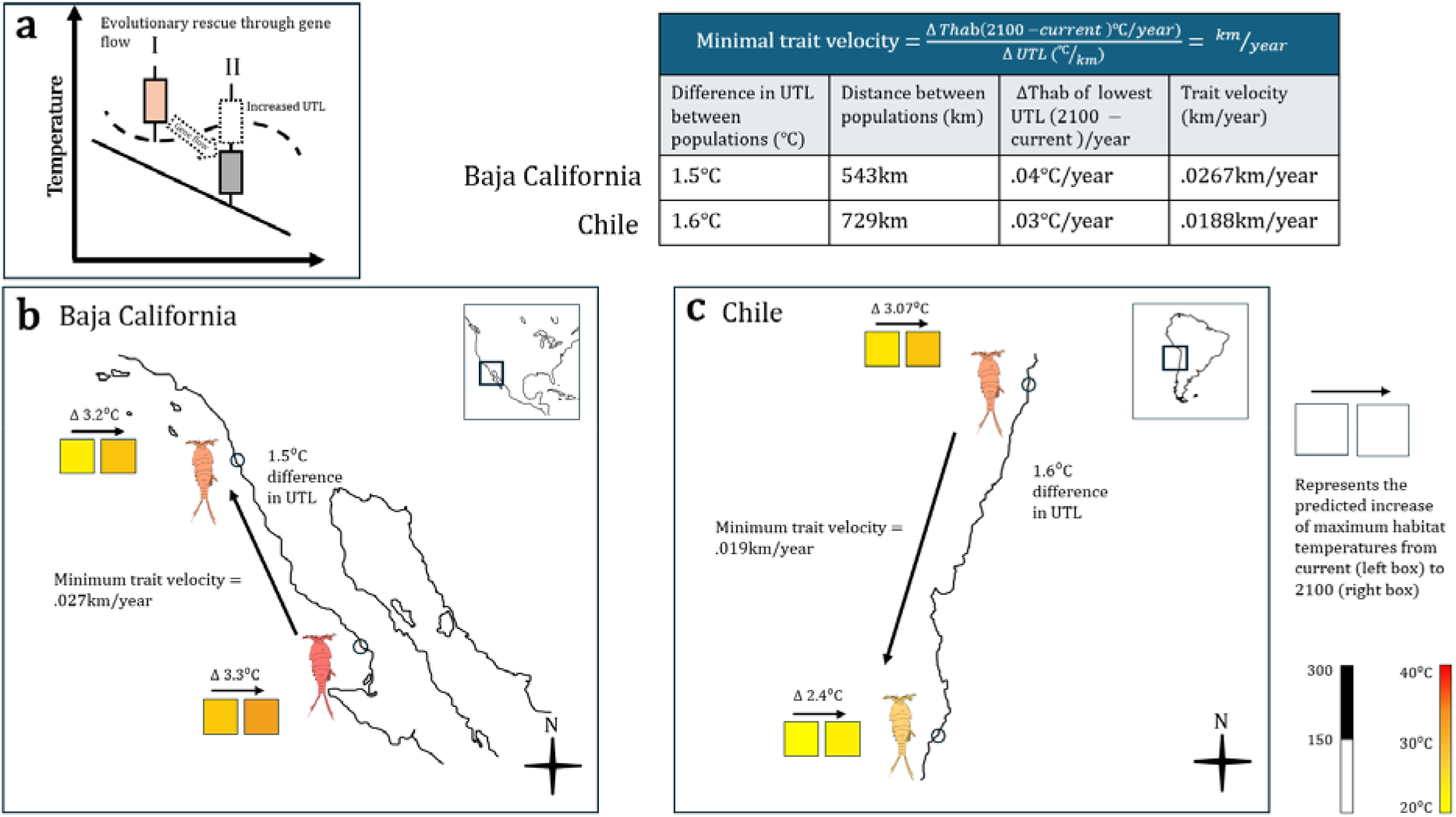

For determining the minimum trait velocity, we incorporate both the variation in thermal tolerance between populations, and the rate of climate warming at each site. To illustrate evolutionary rescue through gene flow, in panel (*a*), population II is expected to experience habitat temperatures above its upper thermal limit (UTL) in the future, resulting in extirpation unless there is evolutionary change in its UTL, which could be facilitated by gene flow from population I, raising UTLs above maximum future temperatures at site II. Panels (c) and (d) present a case study of measuring minimum trait velocity for an intertidal copepod *Tigriopus* sp. along the western coast of North (latitude and longitude 28.65, -114.25 and 32.67, -117.242) and South America (-33.46, -71.67, and -39.86, -73.39), respectively. In both case studies, we calculate the difference in upper thermal limit between the populations (℃), the Euclidian distance (km), and the rate of habitat warming for both populations. The table shows that the potential for evolutionary rescue through gene flow will depend on the rate of warming for the population with the lowest thermal tolerance, the distance between populations, and the difference in thermal tolerance between populations. For example, populations in (b) and (c) have a similar level of intraspecific variation for heat tolerance, but the distance between thermally divergent populations is greater in (c). Still, populations in (c) have a lower minimum trait velocity, because the local rate of warming is lower, reducing the minimum speed of migration from the more tolerant population to the less tolerant population required for rescue (b). All else being equal, the higher latitude population in North America will require a greater minimum trait velocity to keep pace with climate change as opposed to the population from South America. Data from North America based on Kelly et al. (2012). Data From South America are unpublished from Kelly et. al. (in prep.) following methods used in Kelly et al. (2012).

## Materials & Methods

### Population upper thermal limit data

To compare how the distribution of thermal traits across populations varies among species, we used published data on thermal tolerances collected on February 1, 2022, as a part of a previous meta-analysis (Sasaki et al. 2022). Briefly, data was extracted from studies that met several criteria for inclusion: studies measured thermal tolerance limits of ectotherms in temperature (℃) rather than units of time, measured thermal tolerance for two or more populations, measured organism-level thermal tolerance (such as CTmax or LD50); had available data on number of individuals measured from each population; and individuals measured were maintained in common garden conditions across all compared populations. Data were then extracted on the population spatial coordinates, and ecosystem type (i.e. freshwater, intertidal, marine, and terrestrial). For the current analysis, we further filtered the data to focus on the observations reported for upper thermal tolerance and adult life stages only. For a diagram of the entire workflow see supplemental S1.

The original dataset contained individual thermal tolerance measurements, thus for those studies we calculated the mean thermal tolerance of the individuals to obtain population-level means for comparability with studies that reported only population-level measurements. For studies that reported measurements for both males and females, we used the mean of both sexes. We also include and classify the studies according to whether they compared populations across latitudes or elevation. In total we used 79 studies containing 337 populations from 69 different species.

### Population level habitat temperatures and rates of warming

We used two climate databases, Bio-Oracle (version 3.0; Assis et al. 2024) and WorldClim (version 2.1; Fick & Hijmans 2017) to extract present and future climate data from each of the population coordinates in our dataset. For future climate projections from both databases, we used future temperatures projected under the Shared Socioeconomic Pathway (SSP) 4.5 scenario, which represents a ‘middle of the road’ scenario of minimum reduction of greenhouse gas emissions by 2100. Bio-Oracle was used for those populations belonging to marine ecosystems, and we used the sea surface temperature (SST) raster layer that was obtained by averaging data from 10 atmosphere-ocean coupled general circulation models (AO-GCMs, Assis et al. 2024, see Supplemental Figure S1, ST1). WorldClim was used for those populations belonging to freshwater, intertidal and terrestrial ecosystems, and the raster layers corresponded to the air temperature. Air temperature alone may not accurately represent the temperatures that intertidal species experience because body temperature can be influenced by a mixture of atmospheric, oceanic, and biological processes at the local scale (Helmuth et al. 2006, Jawad et al. 2024). However, maximum air temperatures provide a more accurate approximation of the thermal stress that intertidal species can experience compared to sea surface temperature, which does not account for the thermal variability and extremes of emersion.

Bio-Oracle has a spatial resolution of 0.05 degrees (∼5 km) for SSTs. For present climate data, SSTs were calculated using oceanic temperature from maximum values of raster layers for 2000–2010 because data is not available for the current years (2020-2030). For future climatic data, we combined raster layers with maximum SST from 2080–2090 and 2090–2100 to create comparable timeframes to the data available in WorldClim, and we averaged the maximum temperature between layers (Supplemental Figure S1). We extracted maximum monthly SST from Bio-Oracle and extracted air temperature data from Worldclim for the present climatic data (1970–2000) at 2.5 minute (∼5km) spatial resolution using the BIO5 climatic variable which provides values of the ‘maximum temperatures of the warmest month’. For the future climatic data, we used the 2.5 minute spatial resolution for the time period 2081–2100, the same SSPs and BIO5 raster layers. The mean value for each coordinate was extracted by averaging data from raster layers from 6 GCMs that are shared with GCMS used by Bio-Oracle (Supplemental Figure S1, ST1).

### Warming tolerance

For each population we estimated warming tolerance, or the difference between the population’s upper thermal limit and the maximum temperatures of the population’s collection site (Deutsch et al. 2008). This metric has also been referred to as the thermal safety margin (Clusella-Trullas et al. 2021). To account for the effect of acclimation to laboratory temperatures on a population’s thermal tolerance in each study, we calculated a corrected thermal tolerance value following the same methods as Sasaki et al. 2022. Briefly, we first categorized studies based on whether a population’s thermal tolerance was measured from only one acclimation temperature, or if the authors used two or more acclimation temperatures. In the latter case, we used the reaction norm of the thermal tolerance to generate a predicted thermal tolerance based on the mean temperature of the population’s site, what we term in the data set ‘field tolerance’. For studies where only one acclimation temperature was used, we used the reaction norms generated from the above method to create an acclimation response ratio (ARR) for each population based on the global data set. The adjusted thermal tolerance of these populations was predicted using ecosystem type and thermal tolerance from the previous reaction norms as functions for the model (see Sasaki et al. 2022 for complete workflow, see also Pinsky et al. 2019 for alternative ARR method). We then merged the adjusted population upper thermal limit (UTL) data with the average maximum temperatures at the population’s site to calculate the population’s warming tolerance.

We then tested two central questions regarding warming tolerance using these data: First, how does incorporating local adaptation impact estimates of warming tolerance compared to using a species-level thermal tolerance estimates, and does the magnitude of impact differ between ecosystem types? Second, how does incorporating local rates of warming - that is, rates of warming calculated at the populations site, rather than an average rate of warming taken across all populations – impact warming tolerance?

To systematically address these questions, we compared four scenarios based on two binary factors: whether local adaptation (LA) or no local adaptation (NLA) was incorporated in thermal tolerance estimates, and whether local warming (LW) or no local warming (NLW; i.e., average species-level warming) was used in warming rate calculations. The four scenarios are:

1. NLA + NLW: Thermal tolerance is species-level (no local adaptation) and warming rate is averaged across the species (no local warming).
2. NLA + LW: Thermal tolerance is species-level, but warming rate is local to each population.
3. LA + NLW: Thermal tolerance is population-specific (local adaptation), but warming rate is averaged across the species.
4. LA + LW: Both thermal tolerance and warming rate are population-specific (fully local scenario).

Warming tolerance was calculated under current and future conditions for each scenario, allowing us to test the effects of accounting for intraspecific variation in thermal tolerance (LA vs. NLA) and spatial variation in warming rates (LW vs. NLW).

For the NLA scenarios every population within a species was assigned the thermal tolerance of the most tolerant population. This is comparable to estimates of species vulnerability based on climate envelope models, which used the range of conditions tolerated by the species as a proxy for the thermal niche of every population (Angert et al. 2011, Cacciapaglia & van Woesik 2018). We estimated warming tolerance for each population under the NLA scenario and compared the estimates to warming tolerances under the LA scenario, where each population was assigned its own empirically measured thermal tolerance value. For the LW and NLW scenarios we compared warming tolerance estimates of populations calculated with future (2100) temperatures predicted by the local rate of warming at each site (local rate of warming) and compared these to estimates calculated using an average rate of warming across all populations for that species. Furthermore, we compared maximum habitat temperatures between current and future climatic predictions (Supplemental S2, ST3).

### Minimum trait velocity

To examine the potential for evolutionary rescue through gene flow at the pace of climate warming, we compared the spatial distribution of heat tolerance phenotypes across populations to the rate of habitat warming. We used the thermal tolerance values measured for the lowest acclimation temperature in each study to reduce the effects of plasticity for studies that used multiple acclimation temperatures. First, the rate of habitat warming for each population was calculated from the time of the study to the projected average maximum temperatures by 2100. Then, we estimated the minimum trait velocity by dividing the local rate of temperature change (degrees/yr) by the slope of the phenotypic cline in thermal tolerance (degrees of UTL/distance) (see Text Box 1 for visualization).This allowed us to estimate the speed at which the phenotypic isotherm for UTL would need to travel across space to keep pace with rising temperatures, measured in units of km/yr:

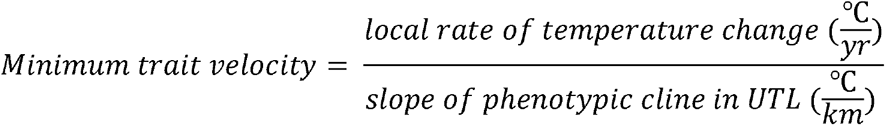

We then measured the Euclidean distance (in km) between each pair of populations using their latitude and longitude by applying the function *distm* (R Core Team, 2025). For studies that did not report the year of collection, we used the year of publication (n studies = 11). Additionally, we used the package *taxize* (Chamverlain and Szöcs, 2013) to add class, order and family of the taxon by obtaining the information from the National Center for Biotechnology Information. Furthermore, we classified the populations by continent using the package *sp* (Pebesma and Bivand, 2005), or by searching their coordinates in Google maps (n populations = 87). We excluded population pairs found on separate continents or oceans, where gene flow is unlikely to occur naturally, which in this data set were populations with a Euclidean distance more than 3730 km. We also excluded population comparisons in which the difference in thermal limits is lower than the per year amount of habitat warming of the population with the lowest UTL, because those comparisons did not contain populations with thermal phenotypes that could provide evolutionary rescue via gene flow given the current rate of warming (pair-wise comparisons excluded = 64). See Supplemental Figure S1 for a full diagram of the workflow. Finally, we assessed the proportion of pair wise population comparisons where the difference in UTL is greater than the projected increase in maximum habitat temperatures in 25 years, as an estimate for the potential for evolutionary rescue via gene flow over the next 25 years.

### Statistical Analysis

We used generalized linear mixed models (GLMMs) to test the influence of incorporating local adaptation on warming tolerance across different ecosystem types. To estimate the effect of local adaptation under current climate conditions, we fitted a GLMM with warming tolerance as the Gaussian-distributed response variable because data were continuous and exhibited positive or negative values. Scenario (with or without local adaptation), ecosystem type, and their interaction were included as fixed effects. Absolute latitude was included as a fixed effect fixed effect to account for its known influence on estimates on warming tolerance (Deutch et al. 2008). To account for non-independence and heterogeneity in the data, we included study and taxon as crossed random effects. Additionally, we incorporated a group-level variance structure scenario and ecosystem using a dispersion parameter (dispformula = ∼ scenario + ecosystem) to model the substantial heteroscedasticity between these groups (Supplemental ST4). Post-hoc pairwise comparisons within groups were conducted using Tukey tests via the *emmeans* package (Lenth, 2021). In general, *glmmTMB* uses asymptotic maximum-likelihood estimation and tests are based on z-statistics with infinite denominator degrees of freedom (*df*, Brooks et al., 2017, Knudson et al 2021), thus *df* are not reported for the post-hoc analysis (Supplemental ST5 & ST7). A second GLMM was fitted with an identical structure to the first but used future warming tolerance estimates to understand how warming tolerance estimates differed between scenarios and ecosystems with and without local adaptation and local rates of warming. Post-hoc Tukey tests were similarly performed to identify significant pairwise differences among ecosystem and scenario groups.

To model minimum trait velocity, we also used generalized linear mixed models. Minimum trait velocity was specified as the Gamma-distributed response variable with a log link function because data was continuous and showed zero or positive values. Ecosystem type was included as the sole fixed effect, with a group-level variance parameter for ecosystem to handle heterogeneous variances (dispformula = ∼ ecosystem). Study and taxon were included as crossed random effects. All models were fitted using the *glmmTMB* package (Brooks et al., 2017) in R version 4.5.2 (R Core Team, 2025), and model assumptions were verified using the *performance* package (Lüdecke et al., 2021). Finally, we evaluated whether existing variation in upper thermal limits (UTL) among populations is sufficient to keep pace with projected rates of warming in the near future. We calculated the proportion of population comparisons in which the difference in UTL was less than the projected increase in maximum habitat temperature expected over the next 25 years from the current data. All data visualization was performed using the *ggplot2* package (Wickham, 2011).

## Results

### Relative rates of climate warming among ecosystem types

Maximum habitat temperatures increased on average by 3.5℃ - 4℃ from study year to 2100 across freshwater, intertidal, marine and terrestrial ecosystems under the future middle-of-the-road SSP2-4.5 climatic scenario (Supplemental S2, ST2). Furthermore, we found that mean rates of habitat warming were similar across ecosystems (freshwater = .045± SE .003; intertidal = .039± SE .001; marine = .043 ± SE .002; terrestrial = .044 ± SE .001 ℃/year; Supplemental ST2). However, the rates of habitat warming varied widely among population sites (0.020 - 0.072 ℃/year), suggesting considerable variation in the rate of warming between populations and species within ecosystems (Supplemental ST3).

### The role of local adaptation in current warming tolerance

Incorporating local adaptation has a strong effect on estimates of warming tolerance for populations in the current era (*χ*^2^ = 62.31, *df* = 1, *p* < 0.001, Fig. 2a-b), while no overall differences in warming tolerance among ecosystems were detected (*χ*^2^ = 5.03, *df* = 3, *p* = 0.17). The effect of incorporating of local adaptation interacted strongly with ecosystem type (scenario x ecosystem interaction: *χ*^2^ = 29.21, *df* = 3, *p <* 0.001) and was associated with larger reductions in mean population-level warming tolerance in marine (*estimate* = - 3.70, *SE* =1.35, *p* = 0.006) and intertidal (*estimate* = -2.78, *SE* = 0.97, *p* = 0.004) ecosystems. In contrast, the reduction in warming tolerance in terrestrial ecosystems (*estimate* = -0.20, *SE* = 0.85, *p* = 0.81) did not differ significantly from that observed in freshwater ecosystems (Supplementary ST4). However, post hoc comparisons revealed significantly lower warming tolerance under local adaptation compared to no local adaptation within intertidal (Δ = −3.74 °C, *z* = 6.94, *p* < 0.001), marine (Δ = −4.57 °C, *z* = 4.28, *p* < 0.001), and terrestrial ecosystems (Δ = −1.21 °C, *z* = 4.88, *p* < 0.001), demonstrating that incorporating local adaptation reduces estimates of warming tolerance across all ecosystem types except for freshwater (Table 2, Fig. 2a-b, Supplementary ST5).

**Table 1.** Mean warming tolerance (WT) estimates (℃) with standard error (SE) for each ecosystem type and the difference in warming tolerance between scenarios. Scenarios reflect those in Figure 2 and methods section under *warming tolerance. NLA no local adaptation, NLW No Local Warming*. Sample sizes (populations): freshwater = 9, intertidal = 60, marine = 33, terrestrial = 210. Mean and standard error (SE) are provided.

| Ecosystem | Scenario | Current<br>mean WT ±<br>SE (°C) | Future<br>mean WT ±<br>SE (°C) |
| --- | --- | --- | --- |
| Freshwater | NLA + NLW | 9.23±1.71 | 5.22±1.91 |
|  | NLA + LW | 9.23±1.71 | 5.22±1.93 |
|  | LA + NLW | 8.35±1.63 | 4.34±1.85 |
|  | LA + LW | 8.35±1.63 | 4.34±1.86 |
| Intertidal | NLA + NLW | 16.78±0.54 | 13.27±0.57 |
|  | NLA + LW | 16.78±0.54 | 13.27±0.53 |
|  | LA + NLW | 13.04±0.50 | 9.53±0.52 |
|  | LA + LW | 13.04±0.5 | 9.53±0.5 |
| Marine | NLA + NLW | 14.64±1.38 | 10.74±1.29 |
|  | NLA + LW | 14.64±1.38 | 10.74±1.2 |
|  | LA + NLW | 10.07±1.84 | 6.16±1.73 |
|  | LA + LW | 10.07±1.84 | 6.16±1.72 |
| Terrestrial | NLA + NLW | 13.47±0.41 | 9.4±0.39 |
|  | NLA + LW | 13.47±0.41 | 9.4±0.39 |
|  | LA + NLW | 12.26±0.41 | 8.19±0.39 |
|  | LA + LW | 12.26±0.41 | 8.19±0.4 |

**Table 2.** Projected minimum trait velocity between all four ecosystems calculated as km/year. Population comparisons were filtered with geographical continuity (less than 3730 km apart), and the difference in UTL of high and low population comparisons is greater than the rate of habitat warming in the lowest population. Mean, standard error (SE), median and number of observations are provided.

| Ecosystem | Mean (km/yr) ± SE | Median (km/yr) | Range (km/yr) | Population contrasts |
| --- | --- | --- | --- | --- |
| Freshwater | 53.06±37.71 | 16.30 | 1.84—390.01 | 10 |
| Intertidal | 63.25±11.36 | 28.77 | 0.11—1814.05 | 241 |
| Marine | 80.47±19.44 | 26.87 | 0.14—1368.81 | 95 |
| Terrestrial | 143.01±25.59 | 28.13 | 0.01—10695.52 | 503 |

**Figure 2.**
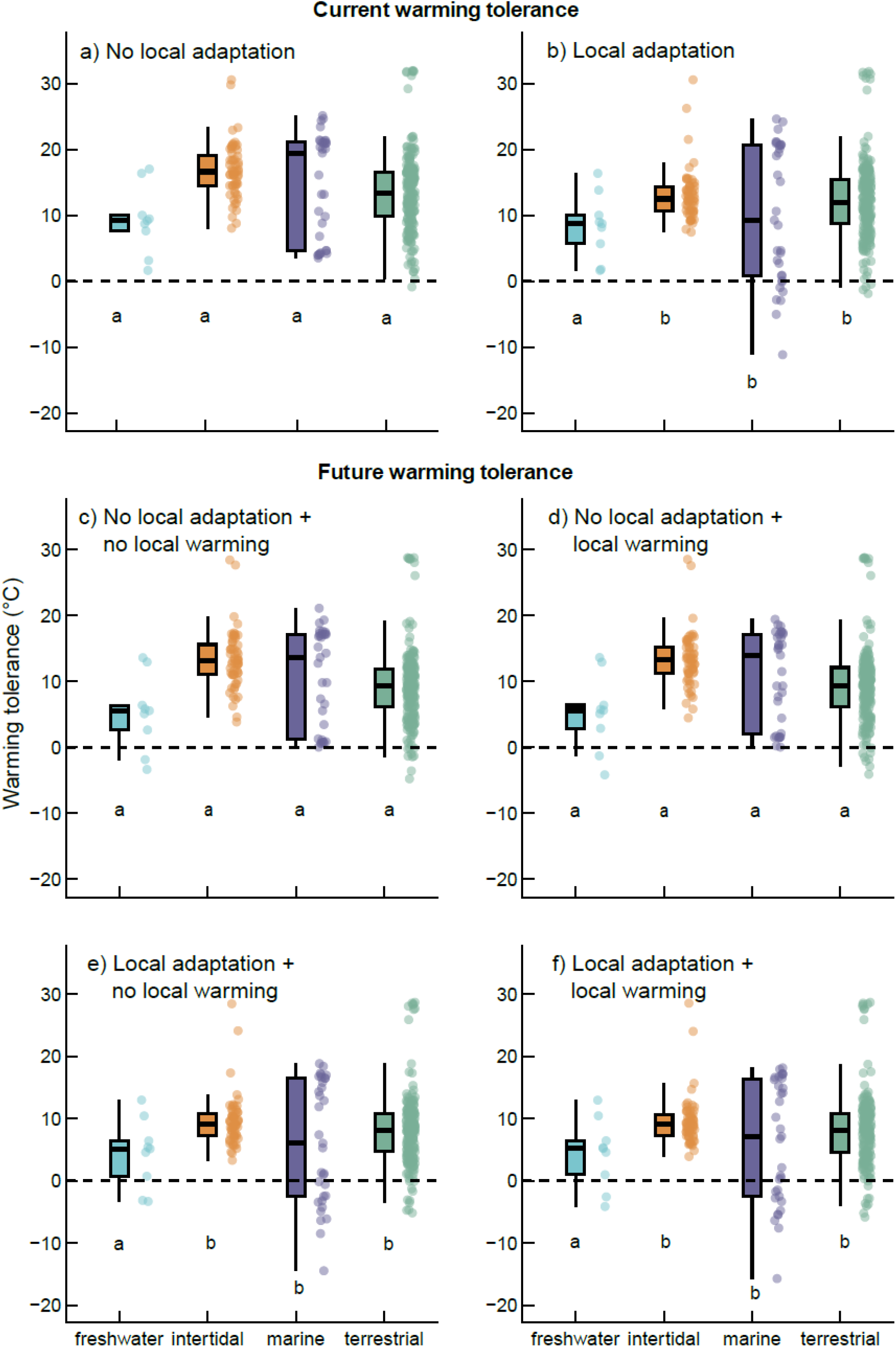
Estimated warming tolerance across ecosystem types both incorporating local adaptation and local rates of warming. Box plots represent the distribution of population-level warming tolerance estimates for each ecosystem in ℃. Letters indicate significant differences among scenarios within ecosystems based on Tukey-adjusted post hoc comparisons (α = 0.05). Warming tolerance in (*a*) was calculated using a standard rate of habitat warming (mean of the maximum habitat temperatures across all populations of a species), without including local adaptation (using the most thermally tolerant populations UTL for all populations, see methods section *Warming tolerance*) in current climate. In (*b*), it was calculated using local rates of habitat warming for each population but with the same shared UTLs across all populations as in (a) again in current climate. In (*c*), future warming tolerance was calculated using the same values as in (*a*). (*d*) incorporates local rates of habitat warming but no local adaptation and (*e*) incorporates local adaptation but without local rates of habitat warming into calculations of future tolerance. Finally, (*f*) calculates future tolerance incorporating both local rates of habitat warming and local adaptation. Boxplots represent the median, 25th and 75th percentiles. Whiskers are values within 1.5× interquartile, black dots represent outliers.

Absolute latitude had a strong positive effect on current warming tolerance (*χ*^*2*^ = 147.69, *df* = 1, *p* < 0.001), with populations at higher latitudes exhibiting higher warming tolerance (supplemental S4, ST4). Residual variance was significantly higher in marine ecosystems (dispersion: z = 4.4, *p* < 0.001), suggesting greater heterogeneity in warming tolerance within this group. Random intercepts indicated substantial among-study variation (variance = 38.35, SD = ±6.19) and moderate among-taxon variation (variance = 5.75, SD = ±2.40; Supplemental ST4).

### The role of local adaptation and local rates of warming on future warming tolerance

Warming tolerance differed strongly among scenarios (*χ*^*2*^ = 128.46, *df* = 3, *p* < 0.001), whereas no overall main effect of ecosystem was detected when averaged across scenarios (*χ*^*2*^ = 5.35, *df* = 3, *p* = 0.15). However, scenario effects depended strongly on ecosystem type, as evidenced by a significant scenario × ecosystem interaction (*χ*^*2*^ = 61.60, *df* = 9, *p* < 0.001). Relative to freshwater systems under the reference scenario A, intertidal ecosystems exhibited higher mean warming tolerance (Supplemental ST6). In contrast, scenarios incorporating local adaptation (scenarios C and D) were associated with pronounced reductions in mean population-level warming tolerance in intertidal and marine ecosystems, whereas corresponding reductions in terrestrial ecosystems were comparatively smaller and not distinguishable from freshwater responses (Supplemental ST6). Incorporating local rates of habitat warming did not significantly impact warming tolerance estimates for any ecosystem type with or without the inclusion of local adaptation (Scenarios A & C *p* > 0.2, ST6).

Post hoc comparisons further supported this pattern, revealing significantly lower warming tolerance under scenarios that incorporate local adaptation (C and D) compared to scenarios without local adaptation (A and B) within intertidal (mean Δ = 3.74), marine (mean Δ = 4.57), and terrestrial ecosystems (mean Δ = 0.91; Table 2, Fig. 2c-f; Supplementary ST7). Variance structure differed among ecosystems, with significantly higher residual variance in marine ecosystems (dispersion: *z* = 6.57, *p* < 0.001) and terrestrial ecosystems (dispersion: *z* = 2.35, *p* = 0.019), indicating greater heterogeneity in warming tolerance within these systems. Random intercepts revealed substantial among-study variation (variance = 38.71, SD = ±6.22) and moderate among-taxon variation (variance = 6.84, SD = ±2.61; Supplemental ST7). Absolute latitude showed a strong positive effect on warming tolerance (*χ*^*2*^ = 256.40, *df* = 1, *p* < 0.001), with populations at higher latitudes exhibiting greater future warming tolerance (Supplementary Fig. S4). Intertidal and terrestrial ecosystems have an increased warming tolerance at higher latitudes, while marine ecosystems appear to have lower warming tolerance at latitudes between 20-30 and 50-60, but highest between 30-40. Freshwater habitats had lower warming tolerance at higher latitudes, but we caution interpreting these results due to limited sampling of freshwater ecosystems at lower latitudes (supplemental S4).

### Minimum trait velocity

Minimum trait velocities varied widely across population comparisons, spanning over five orders of magnitude (0.01–10,695.52 km/year; Table 3). Mixed-effects modeling revealed that mean trait velocity did not differ among ecosystems (χ^2^ = 0.11, df = 3, p = 0.99; Fig. 3a-b). Relative to freshwater ecosystems (reference level), none of the other ecosystem types showed significant differences in model-estimated mean trait velocity, with small and non-significant log-scale effect estimates for intertidal (0.073), marine (0.059), and terrestrial ecosystems (0.212; Supplemental ST8).

**Figure 3.**
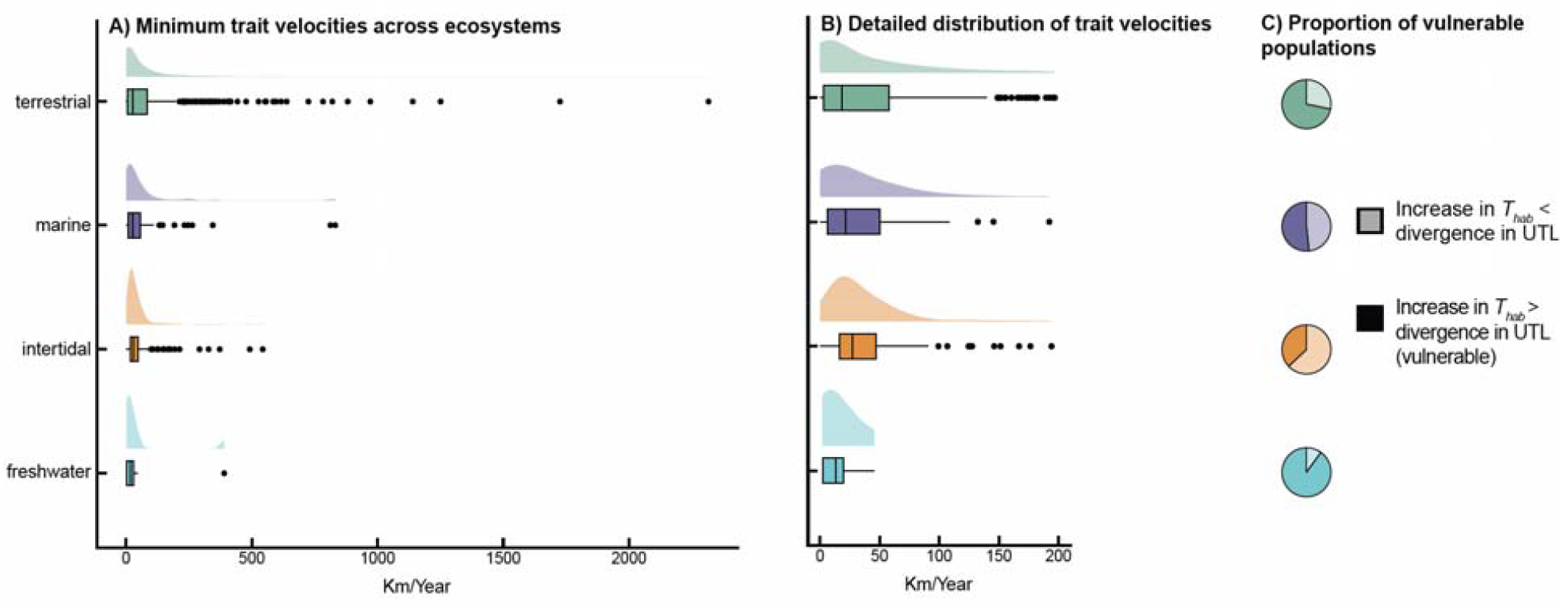
Projected minimum trait velocity and change in habitat temperatures relative to population divergence. (A) Minimum trait velocity across ecosystem types. Calculated values (km/year) for all four ecosystems types from geographically continuous populations pairs where the difference in upper thermal limit (ΔUTL) was sufficient to outpace habitat warming. (B) Distribution of trait velocities, a close-up of (A) to better visualize data density and distribution. (C) relative to divergence, the pie charts display the proportion of population comparisons ∆UTL is less than the predicted habitat temperature increase (∆T_hab_) by near future. Boxplots in (A) and (B) represent the median, 25th and 75th percentiles, and whiskers extending to 1.5× interquartile range; outliers are shown as black dots.

The variance structure did not differ among ecosystems, indicating comparable levels of variability in trait velocity across ecosystem types (Supplementary Table ST8). Random study effects indicated modest among-study variation (variance = 1.54, SD = 1.24) and limited among-taxon variation (variance = 0.29, SD = 0.54), suggesting that differences among studies and taxa contributed relatively little to overall variation in trait velocity (Supplemental ST8). Although summary statistics indicated higher mean trait velocities in terrestrial and marine systems (Table 3), trait velocity distributions were highly right-skewed and characterized by substantial within-ecosystem variability, as reflected by large ranges and medians that were consistently much lower than means. These patterns indicate that apparent differences in trait velocity summaries do not translate into significant differences in ecosystem-level mean trait velocity after accounting for variance structure and random effects (Supplemental ST8). This analysis revealed that most population pairs in freshwater (90%), terrestrial (71%), and marine (51%) ecosystems, as well as 37% of intertidal population pairs, exhibit less variation in upper thermal limits (UTL) than the projected increase in maximum habitat temperature (Fig. 3C; Table S9).

## Discussion

Climate warming is expected to reshape ecosystems by causing local extirpations and range shifts, as populations expand and contract to track suitable habitat temperatures (Chen et al. 2011, Wernberg et al. 2016, Birkett et al. 2018). Comparisons between ecosystem types can provide insight into differences in the effects of warming across broad spatial and taxonomic scales (Jørgensen et al. 2022, Pinsky et al. 2019, Sunday et al. 2019). We demonstrate that using species-level thermal tolerance can underestimate vulnerability across marine, intertidal, and terrestrial ecosystem types, because it assumes that the maximum thermal tolerance across a species range is conserved across populations. Consistent with our predictions, we found that ignoring intraspecific variation in heat tolerance will overestimate warming tolerance the most in marine compared to terrestrial taxa.

Contrary to expectations, incorporating local rates of warming did not impact warming tolerance estimates regardless of whether the scenario incorporated local adaptation. It may be that considering population-specific warming rates increases projected vulnerability for some, while decreasing vulnerability in others, leading to a net effect of zero. However, incorporating fine scale estimates of warming is still likely to be important for population-specific projections of vulnerability, given that rates of warming can differ dramatically between populations within the same species (Supplemental ST3). Indeed, incorporating local rates of warming resulted in greater variance in warming tolerance estimates (Fig. 2), highlighting the role of local-scale climatic processes to shape population-level vulnerability to climate warming (Helmuth et al. 2006). We caution that our estimates of warming tolerance using air surface temperatures may over- or underestimate the realized body temperatures that terrestrial, intertidal, and freshwater taxa could experience, and are used here as a point of comparison between scenarios, but not as predictions of when populations may be at risk of extinction. Body temperatures in these ecosystems can be influenced by a suite of climatic and ecological variables alongside air temperature (Helmuth et al. 2006, Muñoz 2022, Jawad et al. 2024), which will be vital to incorporate in future work aiming to accurately predict the realized exposure to thermal stress in these ecosystems (see Pinsky et al. 2019, Buckley et al. 2015 for incorporating projected body temperatures for estimating vulnerability to warming).

Our results indicate that species with similar geographic ranges may nevertheless be differentially vulnerable to warming. When models of climate-driven range shifts use a species’ geographic range as a proxy for its thermal niche, they assume that every population has the same environmental tolerance according to the highest temperatures found along the species range (Angert et al. 2011, Cacciapaglia & van Woesik 2018), which can overestimate the species projected resilience to habitat warming. These models also tend to project range shifts that occur through extirpation of warm-edge populations (Vilà-Cabrera et al. 2019, Hampe & Petit et al. 2005). However, population-level variation in thermal tolerance, as well as variation in the rate of habitat warming across the species range, will likely result in a mosaic of population level responses to climate warming (Villeneuve et al. 2021, Beaty et al. 2023, Pinsky et al. 2013, Helmuth et al. 2006).

Evolutionary rescue through gene flow is considered a potential mechanism to buffer the negative impacts of habitat warming (Baiotto & Guzman 2025, Brauer et al. 2023, Razgour et al. 2019). Greater divergence in UTL over shorter distances in marine compared to terrestrial taxa would suggest that the potential for evolutionary rescue through gene flow should be greater in marine taxa (Sasaki et al. 2022), but that this will also depend on the relative rates of warming at the populations sites. We evaluated the potential for evolutionary rescue through gene flow, by asking how fast the current phenotypic isoclines within species would have to shift to keep pace with local rates of warming, using the ‘Minimum Trait Velocity’. Surprisingly, despite greater divergence over shorter distances in marine taxa, our results project a similar minimum trait velocity across all four ecosystem types, suggesting that the potential for evolutionary rescue through gene flow could be largely driven by the relative rates of warming at each population’s location rather than by the spatial scale of local adaptation. Indeed, across all ecosystems, most taxa had a limited intraspecific differentiation in UTL relative to the increases in maximum habitat temperatures predicted in the near future (Fig. 3). In other words, the difference in maximum habitat temperature will be greater by the end of the century than the existing differences in phenotypic variation in heat tolerance between populations for most populations of most species, supporting previous findings that potential benefits of evolutionary rescue are highly contingent on lower rates of environmental change (Haley et al. 2013). However, in the near future (25 years from current), marine taxa had more populations with greater divergence between them than the projected increase in habitat temperatures, suggesting greater divergence in marine taxa could still provide some level of buffering against near term warming if such traits are shared between populations as would be expected because of greater dispersal and population connectivity in the ocean. We predict that while gene flow of heat adapted alleles could increase adaptive capacities for populations to near future warming, evolutionary rescue through gene flow has a limited capacity to prevent population extirpation and species range shifts under the full extent of projected changes in maximum temperatures by the end of the century.

There are several important limitations to the methods used here to calculate minimal trait velocity. Our calculations may be overestimates, because they are based only on sampled populations, and do not account for other proximate populations that may be more thermally tolerant. Producing informative minimum trait velocity values for specific species or ecosystems will require increasing sampling across a large or entire portion of a species range to account for more of the variation in thermal tolerance among populations. Here we assume that there is not any systematic bias in population sampling distances across different types of ecosystems. In addition, minimum trait velocity was calculated using Euclidean distance between populations and does not consider variation in biogeographic constraints or facilitators of population connectivity, such as ocean currents or mountain passes (Álvarez-Noriega et al. 2020, Caplat et al. 2016).

For many taxa in each ecosystem, our analysis reveals that variation in heat tolerance between populations is insufficient for evolutionary rescue via adaptive gene flow to keep pace with habitat warming. However, populations with lower heat tolerance could harbor rare variants of heat tolerant alleles, which could facilitate adaptation *in situ*. The presence and frequency of such variants likely depends on the costs of maintaining increased heat tolerance relative to historic maximum habitat temperatures, as well as the amount of gene flow between populations. For example, in *Tigriopus* copepods, individuals with increased heat tolerance can have reduced fecundity, and gene flow between populations is extremely limited (Kelly et al. 2016). As a result, genetic variation in UTLs is limited relative to the variation present at the species level (Kelly et al. 2012). The capacity to evolve higher heat tolerance can also be constrained by biochemical or physiological boundaries to evolving increased heat tolerance at the organismal level (Araújo et al. 2013), and may come at the tradeoff of reduced plasticity in heat tolerance (Barley et al. 2021). Heat tolerance can involve altering a suite of physiological traits and genes, the evolution of which may take considerably longer than the rate of temperature increase species could be experiencing. Recent work has demonstrated that hybrids between thermally divergent populations can result in individuals with more alleles capable of being selected on for increased heat tolerance (Griffiths et al. 2021). Although the capacity to evolve increased thermal tolerance in hybrid individuals could be an important buffer to warming habitats, direct tests of the heat tolerance capacity of hybrid individuals remain scarce, and thus difficult to predict the realized heat tolerance potential of populations that receive more thermally tolerant phenotypes from neighboring populations.

As habitats warm, species thermal phenotypes are increasingly mismatched with their local habitat temperatures. Our study emphasizes the important role of incorporating local adaptation in our predictions of population and species vulnerability to warming across ecosystems. Minimum trait velocity can be a useful predictive metric for determining the potential for evolutionary rescue through gene flow to provide more thermally tolerant alleles at the pace of habitat warming. Assisted gene flow to provide more adaptive genotypes to populations threatened with environmental degradation is becoming increasingly considered as populations of ecologically important species are being extirpated at unprecedented rates (Baker et al. 2025). For systems in which the potential for such intervention is being considered, minimal trait velocity can provide an important predictive metric to assess the spatial scale of phenotypic variation in thermal traits for a species. By incorporating more fine scale details, such as biogeographic barriers and species mobility and known gene flow between populations, the potential for evolutionary rescue through gene flow to occur without intervention can be predicted at the pace of climate change, to determine if populations could be rescued naturally, or if rates of warming exceed the potential for evolutionary rescue. Overall, our study demonstrates that in the near future, the middle-of-the road SSP2-4.5 projection of climate warming will outpace the potential for adaptation based on the standing adaptive differentiation for heat tolerance between populations, suggesting that natural gene flow between populations has limited potential for rescuing populations at the pace of current climate warming. Our results emphasize several key conservation measures to reduce population extirpations as the climate warms. For marine taxa, the creation and maintenance of marine protected areas and networks can help promote gene flow between populations facing rapid environmental change (Van Der Meer et al. 2015, Pazmiño et al. 2017). For intertidal and terrestrial taxa, maintaining habitat heterogeneity through protection of coastal and terrestrial habitats can promote the availability of thermal refugia (Morelli et al. 2016). Finally, we emphasize that efforts on reducing rates of global climate warming will be the most effective action for long term protection against population and species extinctions.

## Acknowledgments

We are grateful to our colleagues at Louisiana State University and beyond for insightful conversations about the many ways species are responding to a warming world. We thank Pablo Moreno Garcia for helping us develop the script used for preparing part of the data set in this study. We thank Edwin Torres for help in identifying suitable future climate projections. We thank Michael E. Hellberg and Kyle E. Harms for helpful suggestions throughout the process of developing this manuscript through its final draft. We thank Lisa M. Komoroske for insightful comments that led to the naming of the term ‘minimal trait velocity’. A.L.S was supported by the National Science Foundation grant DBE 2216631.

## Notes

### Competing Interest Statement

The authors have declared no competing interest.

