## Supplemental material for "A mosaic of climate vulnerability: local warming rates meet intraspecific divergence in heat tolerance"

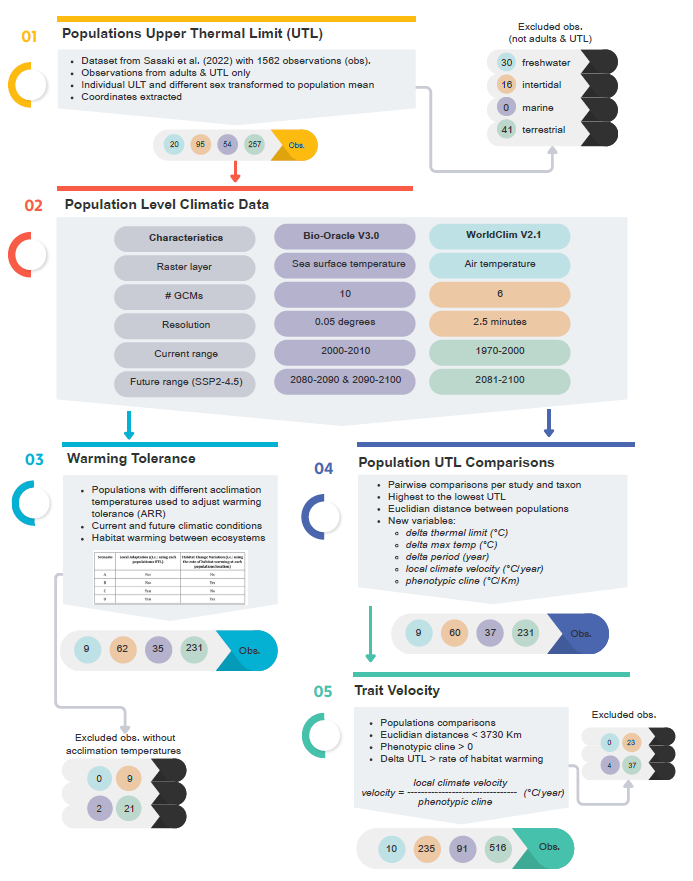

S1 General workflow of data. (1) We started with selecting the data of interest from Sasaki et al. (2022), in total we obtained 337 populations of interest, while a total of 528 were not included because did not corresponded to adults or UTL. (2) Then we obtained climatic data from the populations coordinates of interest using Bio-Oracle for marine (purple) and WorldClim for freshwater (blue), intertidal (orange), terrestrial (green) populations. The characteristics of these datasets were matched as close as possible to avoid particular differences. (3) We created different scenarios that included or not local adaptation or local climate velocity to understand the role of these on warming tolerance of populations. (4) We calculated population UTL comparisons using the rate of local climate velocity and phenotypic cline. (5) With the population UTL comparisons, we calculated the minimum trait velocity with 852 populations comparisons, while 64 were excluded from the analysis because the distance between populations were higher than 3730 km and cannot reach other naturally, and they did not have a difference in the UTL limit higher to their rate of habitat warming.

ST1 Table of shared general circulation models (GCMs) between WorldClim and Bio-Oracle for future projected temperature data. GMC with an asterisk represented a shared model but was not used in the analysis as climatic data for the years of interest was not available.

| **Bio-Oracle V3** | **WorldClim V2.1** |
| --- | --- |
| ACCESS-ESM1-5 108 | ACCESS-CM2 |
| CanESM5 |  |
| CESM2-WACCM 110 |  |
| CNRM-ESM2-1 |  |
| GFDL-ESM4 | GFDL-ESM4* |
| GISS-E2-1-G | GISS-E2-1-G |
| IPSL-CM6A-LR | IPSL-CM6A-LR |
| MIROC-ES2L | MIROCC6 |
| MRI117 ESM2-0 | MRI-ESM2-0 |
| UKESM1-0-LL | UKESM1-0-LL |

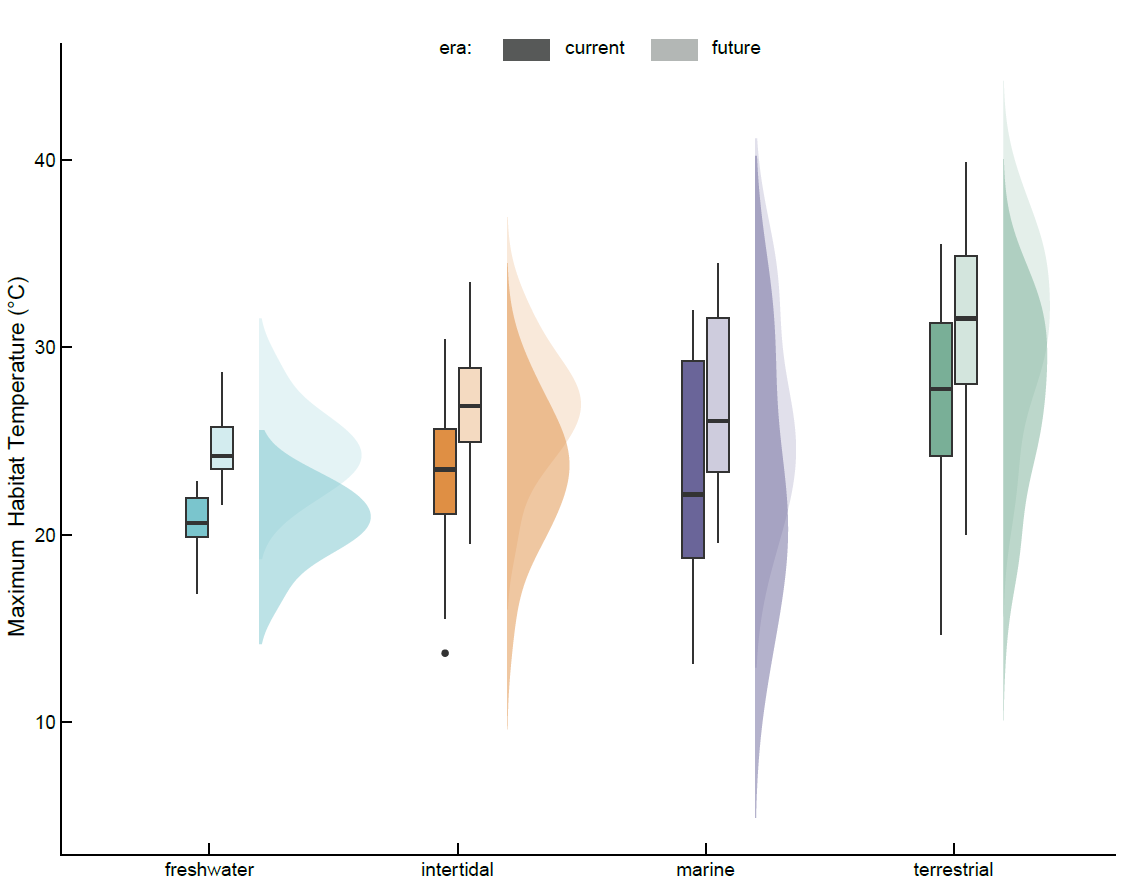

S2 Mean maximum habitat temperatures in each realm. Dark shade is current temperatures; light shade corresponds to future conditions. Box plots represent the median, 25th and 75th percentiles, and whiskers extending to 1.5× interquartile range; outliers are shown as black dots.

ST2 Descriptive statistics of maximum habitat temperatures (°C) across ecosystems for current conditions and future climate projections. Values shown include the mean of maximum temperatures (mean max temp), the minimum of maximum temperatures (lowest value of max temp observed), and the maximum of maximum temperatures (highest value of max temp observed). The number of observations (n pops) per ecosystem is given.

| **Ecosystem type** | **n pops** | **Current conditions** | | | **Future Condition** | | |
| --- | --- | --- | --- | --- | --- | --- | --- |
|  |  | **Mean max temp.** | **Min of max temp.** | **Max of max temp.** | **Mean max temp.** | **Min of max temp.** | **Max of max temp.** |
| Freshwater | 9 | 20.63 | 16.87 | 22.87 | 24.63 | 21.60 | 28.65 |
| Intertidal | 60 | 23.37 | 13.68 | 30.44 | 26.88 | 19.51 | 33.47 |
| Marine | 33 | 22.83 | 13.11 | 31.99 | 26.74 | 19.58 | 34.49 |
| Terrestrial | 210 | 27.18 | 14.65 | 35.50 | 31.25 | 20.00 | 39.87 |

ST3 Summary of rate of habitat warming across ecosystems. Values include number of comparisons (n), average rate (avg), median rate (median), standard error of the mean (SE), and lower and upper bounds of the range. All rates are given in °C.

| **Ecosystem** | **n** | **Avg** | **Median** | **SE** | **Low** | **High** |
| --- | --- | --- | --- | --- | --- | --- |
| Freshwater | 10 | 0.046 | 0.040 | 0.003 | 0.037 | 0.064 |
| Intertidal | 235 | 0.041 | 0.039 | 0.000 | 0.024 | 0.060 |
| Marine | 91 | 0.046 | 0.048 | 0.001 | 0.020 | 0.065 |
| Terrestrial | 485 | 0.039 | 0.031 | 0.001 | 0.023 | 0.072 |

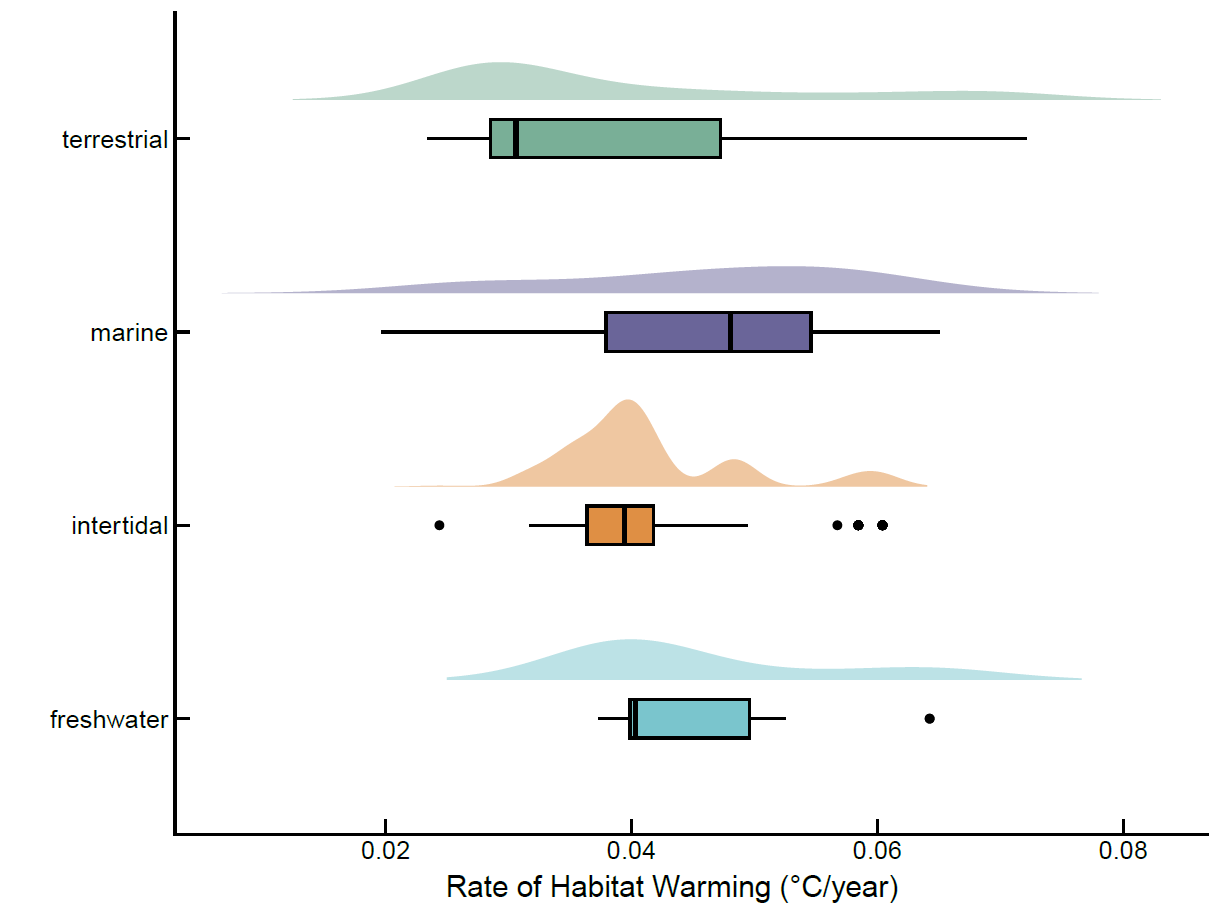

S3 Rate of habitat warming (°C/year) across four ecosystems. Box plots represent the median, 25th and 75th percentiles, and whiskers extending to 1.5× interquartile range; outliers are shown as black dots.

ST4. Parameter estimates from a Gaussian generalized linear mixed-effects model (log link) evaluating the effects of local adaptation scenario (scenario C vs. scenario A), ecosystem type, their interaction, and absolute latitude on current warming tolerance. Fixed-effect estimates are shown for the conditional (mean) model, with freshwater ecosystem and the non-local-adaptation scenario (A) as reference levels. The dispersion model reports effects of scenario and ecosystem on residual variance. Random effects represent among-study and among-taxon variation in intercepts (n = 624, study = 35, taxon = 67). Estimates are reported with standard errors, Wald z-values, and associated p-values. Significant p-values in bold.

| **Effect** | **Group** | **Term** | **Estimate** | **Standard Error** | **Z -value** | **P-value** |
| --- | --- | --- | --- | --- | --- | --- |
| *Conditional Model:* | |  |  |  |  |  |
| fixed | NA | (Intercept) | -1.828 | 4.001 | -0.457 | 0.648 |
| fixed | NA | scenarioC | -0.883 | 0.818 | -1.080 | 0.280 |
| fixed | NA | ecosystemintertidal | 12.030 | 4.765 | 2.524 | **0.012** |
| fixed | NA | ecosystemmarine | 6.354 | 4.857 | 1.308 | 0.191 |
| fixed | NA | ecosystemterrestrial | 6.235 | 4.199 | 1.485 | 0.138 |
| fixed | NA | abs_lat | 0.241 | 0.020 | 12.153 | **0.000** |
| fixed | NA | scenarioC:ecosystemintertidal | -2.784 | 0.973 | -2.861 | **0.004** |
| fixed | NA | scenarioC:ecosystemmarine | -3.703 | 1.348 | -2.746 | **0.006** |
| fixed | NA | scenarioC:ecosystemterrestrial | -0.199 | 0.847 | -0.235 | 0.814 |
| *Dispersion Model:* | |  |  |  |  |  |
| dispersion | NA | (Intercept) | 0.587 | 0.187 | 3.137 | **0.002** |
| dispersion | NA | scenarioC | -0.076 | 0.063 | -1.205 | 0.228 |
| dispersion | NA | ecosystemintertidal | 0.210 | 0.197 | 1.066 | 0.286 |
| dispersion | NA | ecosystemmarine | 0.907 | 0.206 | 4.405 | **0.000** |
| dispersion | NA | ecosystemterrestrial | 0.256 | 0.188 | 1.366 | 0.172 |
| *Conditional Model:* | |  | **Variance** | **Standard Deviation** | |  |
| random | study | sd__(Intercept) | 38.347 | 6.192 |  |  |
| random | taxon | sd__(Intercept) | 5.746 | 2.397 |  |  |
| random | Residual | sd__Observation | NA | NA |  |  |

ST5. Tukey-adjusted pairwise comparisons of model-estimated mean warming tolerance across combinations of local adaptation scenario (A = no local adaptation, C = local adaptation) and ecosystem type, based on estimated marginal means from the mixed-effects model. Estimates represent differences in mean waring tolerance on the response scale between contrast levels (first minus second). Standard errors, Wald z-ratios, and p-values are reported. Freshwater ecosystems and scenario A serve as reference levels in the underlying model. Degrees of freedom are asymptotic and equal to infinite (see methods for details). P-values in bold are significant.

| **ID** | **Contrast** | **Estimate** | **Standard Error** | **Z-ratio** | **P-value** |
| --- | --- | --- | --- | --- | --- |
| 1 | A freshwater - C freshwater | 0.88 | 0.82 | 1.08 | 0.96 |
| 2 | A freshwater - A intertidal | -12.03 | 4.77 | -2.52 | 0.18 |
| 3 | A freshwater - C intertidal | -8.36 | 4.75 | -1.76 | 0.65 |
| 4 | A freshwater - A marine | -6.35 | 4.86 | -1.31 | 0.90 |
| 5 | A freshwater - C marine | -1.77 | 4.83 | -0.37 | 1.00 |
| 6 | A freshwater - A terrestrial | -6.24 | 4.20 | -1.48 | 0.82 |
| 7 | A freshwater - C terrestrial | -5.15 | 4.20 | -1.23 | 0.92 |
| 8 | C freshwater - A intertidal | -12.91 | 4.76 | -2.71 | 0.12 |
| 9 | C freshwater - C intertidal | -9.25 | 4.74 | -1.95 | 0.52 |
| 10 | C freshwater - A marine | -7.24 | 4.85 | -1.49 | 0.81 |
| 11 | C freshwater - C marine | -2.65 | 4.83 | -0.55 | 1.00 |
| 12 | C freshwater - A terrestrial | -7.12 | 4.19 | -1.70 | 0.69 |
| 13 | C freshwater - C terrestrial | -6.04 | 4.19 | -1.44 | 0.84 |
| 14 | A intertidal - C intertidal | 3.67 | 0.53 | 6.94 | **0.00** |
| 15 | A intertidal - A marine | 5.68 | 4.01 | 1.42 | 0.85 |
| 16 | A intertidal - C marine | 10.26 | 3.98 | 2.58 | 0.16 |
| 17 | A intertidal - A terrestrial | 5.79 | 3.17 | 1.83 | 0.60 |
| 18 | A intertidal - C terrestrial | 6.88 | 3.17 | 2.17 | 0.37 |
| 19 | C intertidal - A marine | 2.01 | 3.99 | 0.50 | 1.00 |
| 20 | C intertidal - C marine | 6.59 | 3.96 | 1.67 | 0.71 |
| 21 | C intertidal - A terrestrial | 2.13 | 3.15 | 0.68 | 1.00 |
| 22 | C intertidal - C terrestrial | 3.21 | 3.15 | 1.02 | 0.97 |
| 23 | A marine - C marine | 4.59 | 1.07 | 4.28 | **0.00** |
| 24 | A marine - A terrestrial | 0.12 | 3.31 | 0.04 | 1.00 |
| 25 | A marine - C terrestrial | 1.20 | 3.31 | 0.36 | 1.00 |
| 26 | C marine - A terrestrial | -4.47 | 3.28 | -1.36 | 0.87 |
| 27 | C marine - C terrestrial | -3.39 | 3.28 | -1.03 | 0.97 |
| 28 | A terrestrial - C terrestrial | 1.08 | 0.22 | 4.88 | **0.00** |

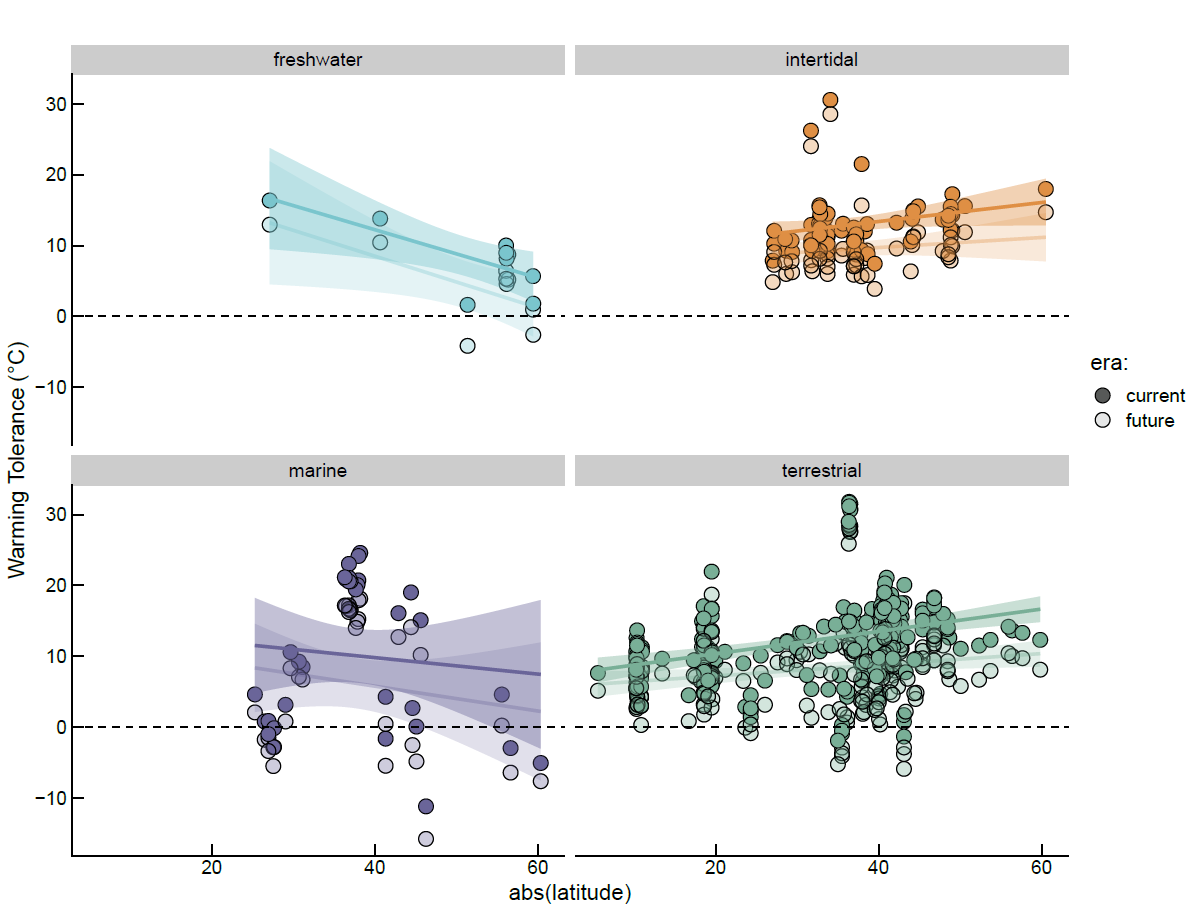

S4 Comparison of warming tolerance across latitudes for each realm in current and 2100. Warming tolerance here included both local adaptation and local climate velocity (scenario D).

ST6. Parameter estimates from a Gaussian generalized linear mixed-effects model (log link) evaluating the effects of local adaptation and local habitat warming, ecosystem type, their interaction, and absolute latitude on future warming tolerance. Fixed-effect estimates are shown for the conditional (mean) model, with freshwater ecosystem and the non-local-adaptation scenario (A) as reference levels. The dispersion model reports effects of scenario and ecosystem on residual variance. Random effects represent among-study and among-taxon variation in intercepts (n = 1248, study = 35, taxon = 67). Estimates are reported with standard errors, Wald z-values, and associated p-values. Significant p-values are in bold.

| **Effect** | **Group** | **Term** | **Estimate** | **Standard Error** | **Z-value** | **P-value** |
| --- | --- | --- | --- | --- | --- | --- |
| *Conditional Model* | |  |  |  |  |  |
| fixed | NA | (Intercept) | -4.873 | 3.997 | -1.219 | 0.223 |
| fixed | NA | scenarioB | 0.000 | 0.800 | 0.000 | 1.000 |
| fixed | NA | scenarioC | -0.883 | 0.767 | -1.151 | 0.250 |
| fixed | NA | scenarioD | -0.883 | 0.788 | -1.121 | 0.262 |
| fixed | NA | ecosystemintertidal | 12.410 | 4.851 | 2.558 | 0.011 |
| fixed | NA | ecosystemmarine | 6.181 | 4.913 | 1.258 | 0.208 |
| fixed | NA | ecosystemterrestrial | 6.196 | 4.252 | 1.457 | 0.145 |
| fixed | NA | abs_lat | 0.222 | 0.014 | 16.012 | **0.000** |
| fixed | NA | scenarioB:ecosystemintertidal | 0.000 | 0.892 | 0.000 | 1.000 |
| fixed | NA | scenarioC:ecosystemintertidal | -2.773 | 0.891 | -3.113 | **0.002** |
| fixed | NA | scenarioD:ecosystemintertidal | -2.773 | 0.913 | -3.036 | **0.002** |
| fixed | NA | scenarioB:ecosystemmarine | 0.000 | 1.312 | 0.000 | 1.000 |
| fixed | NA | scenarioC:ecosystemmarine | -3.679 | 1.264 | -2.910 | **0.004** |
| fixed | NA | scenarioD:ecosystemmarine | -3.679 | 1.299 | -2.833 | **0.005** |
| fixed | NA | scenarioB:ecosystemterrestrial | 0.000 | 0.830 | 0.000 | 1.000 |
| fixed | NA | scenarioC:ecosystemterrestrial | -0.186 | 0.797 | -0.234 | 0.815 |
| fixed | NA | scenarioD:ecosystemterrestrial | -0.186 | 0.819 | -0.228 | 0.820 |
| *Dispersion Model* | |  |  |  |  |  |
| Dispersion | NA | (Intercept) | 0.537 | 0.129 | 4.153 | **0.000** |
| Dispersion | NA | scenarioB | -0.016 | 0.057 | -0.287 | 0.774 |
| Dispersion | NA | scenarioC | -0.105 | 0.060 | -1.753 | 0.080 |
| Dispersion | NA | scenarioD | -0.047 | 0.060 | -0.773 | 0.440 |
| Dispersion | NA | ecosystemintertidal | 0.243 | 0.133 | 1.833 | 0.067 |
| Dispersion | NA | ecosystemmarine | 0.912 | 0.139 | 6.565 | **0.000** |
| Dispersion | NA | ecosystemterrestrial | 0.296 | 0.126 | 2.349 | **0.019** |
| *Conditional Model* | |  | **Variance** | **Standard Deviation** | |  |
| random effect | study | sd__(Intercept) | 38.714 | 6.222 |  |  |
| random effect | taxon | sd__(Intercept) | 6.835 | 2.614 |  |  |
| random effect | Residual | sd__Observation | NA | NA |  |  |

ST7. Tukey-adjusted pairwise comparisons of model-estimated mean warming tolerance across combinations of local adaptation, local climate warming, and ecosystem type, based on estimated marginal means from the mixed-effects model. Estimates represent differences in mean waring tolerance on the response scale between contrast levels (first minus second). Standard errors, Wald z-ratios, and p-values are reported. Freshwater ecosystems and scenario A serve as reference levels in the underlying model. Degrees of freedom are asymptotic and equal to infinite (see methods for details). P-values in bold are significant.

| **ID** | **Contrast** | **Estimate** | **Standard Error** | **Z-ratio** | **P-value** |
| --- | --- | --- | --- | --- | --- |
| 1 | A freshwater - B freshwater | 0.00 | 0.80 | 0.00 | 1.00 |
| 2 | A freshwater - C freshwater | 0.88 | 0.77 | 1.15 | 1.00 |
| 3 | A freshwater - D freshwater | 0.88 | 0.79 | 1.12 | 1.00 |
| 4 | A freshwater - A intertidal | -12.41 | 4.85 | -2.56 | 0.43 |
| 5 | A freshwater - B intertidal | -12.41 | 4.85 | -2.56 | 0.43 |
| 6 | A freshwater - C intertidal | -8.75 | 4.84 | -1.81 | 0.91 |
| 7 | A freshwater - D intertidal | -8.75 | 4.84 | -1.81 | 0.91 |
| 8 | A freshwater - A marine | -6.18 | 4.91 | -1.26 | 1.00 |
| 9 | A freshwater - B marine | -6.18 | 4.91 | -1.26 | 1.00 |
| 10 | A freshwater - C marine | -1.62 | 4.90 | -0.33 | 1.00 |
| 11 | A freshwater - D marine | -1.62 | 4.90 | -0.33 | 1.00 |
| 12 | A freshwater - A terrestrial | -6.20 | 4.25 | -1.46 | 0.99 |
| 13 | A freshwater - B terrestrial | -6.20 | 4.25 | -1.46 | 0.99 |
| 14 | A freshwater - C terrestrial | -5.13 | 4.25 | -1.21 | 1.00 |
| 15 | A freshwater - D terrestrial | -5.13 | 4.25 | -1.21 | 1.00 |
| 16 | B freshwater - C freshwater | 0.88 | 0.76 | 1.16 | 1.00 |
| 17 | B freshwater - D freshwater | 0.88 | 0.78 | 1.13 | 1.00 |
| 18 | B freshwater - A intertidal | -12.41 | 4.85 | -2.56 | 0.43 |
| 19 | B freshwater - B intertidal | -12.41 | 4.85 | -2.56 | 0.43 |
| 20 | B freshwater - C intertidal | -8.75 | 4.84 | -1.81 | 0.91 |
| 21 | B freshwater - D intertidal | -8.75 | 4.84 | -1.81 | 0.91 |
| 22 | B freshwater - A marine | -6.18 | 4.91 | -1.26 | 1.00 |
| 23 | B freshwater - B marine | -6.18 | 4.91 | -1.26 | 1.00 |
| 24 | B freshwater - C marine | -1.62 | 4.89 | -0.33 | 1.00 |
| 25 | B freshwater - D marine | -1.62 | 4.90 | -0.33 | 1.00 |
| 26 | B freshwater - A terrestrial | -6.20 | 4.25 | -1.46 | 0.99 |
| 27 | B freshwater - B terrestrial | -6.20 | 4.25 | -1.46 | 0.99 |
| 28 | B freshwater - C terrestrial | -5.13 | 4.25 | -1.21 | 1.00 |
| 29 | B freshwater - D terrestrial | -5.13 | 4.25 | -1.21 | 1.00 |
| 30 | C freshwater - D freshwater | 0.00 | 0.75 | 0.00 | 1.00 |
| 31 | C freshwater - A intertidal | -13.29 | 4.84 | -2.74 | 0.30 |
| 32 | C freshwater - B intertidal | -13.29 | 4.84 | -2.74 | 0.30 |
| 33 | C freshwater - C intertidal | -9.64 | 4.84 | -1.99 | 0.83 |
| 34 | C freshwater - D intertidal | -9.64 | 4.84 | -1.99 | 0.83 |
| 35 | C freshwater - A marine | -7.06 | 4.91 | -1.44 | 0.99 |
| 36 | C freshwater - B marine | -7.06 | 4.90 | -1.44 | 0.99 |
| 37 | C freshwater - C marine | -2.50 | 4.89 | -0.51 | 1.00 |
| 38 | C freshwater - D marine | -2.50 | 4.89 | -0.51 | 1.00 |
| 39 | C freshwater - A terrestrial | -7.08 | 4.25 | -1.67 | 0.96 |
| 40 | C freshwater - B terrestrial | -7.08 | 4.25 | -1.67 | 0.96 |
| 41 | C freshwater - C terrestrial | -6.01 | 4.24 | -1.42 | 0.99 |
| 42 | C freshwater - D terrestrial | -6.01 | 4.24 | -1.42 | 0.99 |
| 43 | D freshwater - A intertidal | -13.29 | 4.85 | -2.74 | 0.30 |
| 44 | D freshwater - B intertidal | -13.29 | 4.85 | -2.74 | 0.30 |
| 45 | D freshwater - C intertidal | -9.64 | 4.84 | -1.99 | 0.83 |
| 46 | D freshwater - D intertidal | -9.64 | 4.84 | -1.99 | 0.83 |
| 47 | D freshwater - A marine | -7.06 | 4.91 | -1.44 | 0.99 |
| 48 | D freshwater - B marine | -7.06 | 4.91 | -1.44 | 0.99 |
| 49 | D freshwater - C marine | -2.50 | 4.89 | -0.51 | 1.00 |
| 50 | D freshwater - D marine | -2.50 | 4.90 | -0.51 | 1.00 |
| 51 | D freshwater - A terrestrial | -7.08 | 4.25 | -1.67 | 0.96 |
| 52 | D freshwater - B terrestrial | -7.08 | 4.25 | -1.67 | 0.96 |
| 53 | D freshwater - C terrestrial | -6.01 | 4.25 | -1.41 | 0.99 |
| 54 | D freshwater - D terrestrial | -6.01 | 4.25 | -1.41 | 0.99 |
| 55 | A intertidal - B intertidal | 0.00 | 0.40 | 0.00 | 1.00 |
| 56 | A intertidal - C intertidal | 3.66 | 0.45 | 8.08 | **0.00** |
| 57 | A intertidal - D intertidal | 3.66 | 0.46 | 7.93 | **0.00** |
| 58 | A intertidal - A marine | 6.23 | 4.07 | 1.53 | 0.98 |
| 59 | A intertidal - B marine | 6.23 | 4.07 | 1.53 | 0.98 |
| 60 | A intertidal - C marine | 10.79 | 4.05 | 2.67 | 0.35 |
| 61 | A intertidal - D marine | 10.79 | 4.06 | 2.66 | 0.36 |
| 62 | A intertidal - A terrestrial | 6.21 | 3.24 | 1.92 | 0.87 |
| 63 | A intertidal - B terrestrial | 6.21 | 3.24 | 1.92 | 0.87 |
| 64 | A intertidal - C terrestrial | 7.28 | 3.24 | 2.25 | 0.66 |
| 65 | A intertidal - D terrestrial | 7.28 | 3.24 | 2.25 | 0.66 |
| 66 | B intertidal - C intertidal | 3.66 | 0.45 | 8.13 | **0.00** |
| 67 | B intertidal - D intertidal | 3.66 | 0.46 | 7.97 | **0.00** |
| 68 | B intertidal - A marine | 6.23 | 4.07 | 1.53 | 0.98 |
| 69 | B intertidal - B marine | 6.23 | 4.07 | 1.53 | 0.98 |
| 70 | B intertidal - C marine | 10.79 | 4.05 | 2.67 | 0.35 |
| 71 | B intertidal - D marine | 10.79 | 4.06 | 2.66 | 0.36 |
| 72 | B intertidal - A terrestrial | 6.21 | 3.24 | 1.92 | 0.87 |
| 73 | B intertidal - B terrestrial | 6.21 | 3.24 | 1.92 | 0.87 |
| 74 | B intertidal - C terrestrial | 7.28 | 3.24 | 2.25 | 0.66 |
| 75 | B intertidal - D terrestrial | 7.28 | 3.24 | 2.25 | 0.66 |
| 76 | C intertidal - D intertidal | 0.00 | 0.37 | 0.00 | 1.00 |
| 77 | C intertidal - A marine | 2.57 | 4.06 | 0.63 | 1.00 |
| 78 | C intertidal - B marine | 2.57 | 4.06 | 0.63 | 1.00 |
| 79 | C intertidal - C marine | 7.14 | 4.04 | 1.77 | 0.93 |
| 80 | C intertidal - D marine | 7.14 | 4.05 | 1.76 | 0.93 |
| 81 | C intertidal - A terrestrial | 2.56 | 3.23 | 0.79 | 1.00 |
| 82 | C intertidal - B terrestrial | 2.56 | 3.23 | 0.79 | 1.00 |
| 83 | C intertidal - C terrestrial | 3.63 | 3.23 | 1.12 | 1.00 |
| 84 | C intertidal - D terrestrial | 3.63 | 3.23 | 1.12 | 1.00 |
| 85 | D intertidal - A marine | 2.57 | 4.06 | 0.63 | 1.00 |
| 86 | D intertidal - B marine | 2.57 | 4.06 | 0.63 | 1.00 |
| 87 | D intertidal - C marine | 7.14 | 4.04 | 1.77 | 0.93 |
| 88 | D intertidal - D marine | 7.14 | 4.05 | 1.76 | 0.93 |
| 89 | D intertidal - A terrestrial | 2.56 | 3.23 | 0.79 | 1.00 |
| 90 | D intertidal - B terrestrial | 2.56 | 3.23 | 0.79 | 1.00 |
| 91 | D intertidal - C terrestrial | 3.63 | 3.23 | 1.12 | 1.00 |
| 92 | D intertidal - D terrestrial | 3.63 | 3.23 | 1.12 | 1.00 |
| 93 | A marine - B marine | 0.00 | 1.04 | 0.00 | 1.00 |
| 94 | A marine - C marine | 4.56 | 1.01 | 4.54 | **0.00** |
| 95 | A marine - D marine | 4.56 | 1.03 | 4.42 | **0.00** |
| 96 | A marine - A terrestrial | -0.01 | 3.34 | 0.00 | 1.00 |
| 97 | A marine - B terrestrial | -0.01 | 3.34 | 0.00 | 1.00 |
| 98 | A marine - C terrestrial | 1.05 | 3.34 | 0.32 | 1.00 |
| 99 | A marine - D terrestrial | 1.05 | 3.34 | 0.32 | 1.00 |
| 100 | B marine - C marine | 4.56 | 1.00 | 4.58 | **0.00** |
| 101 | B marine - D marine | 4.56 | 1.02 | 4.46 | **0.00** |
| 102 | B marine - A terrestrial | -0.01 | 3.33 | 0.00 | 1.00 |
| 103 | B marine - B terrestrial | -0.01 | 3.33 | 0.00 | 1.00 |
| 104 | B marine - C terrestrial | 1.05 | 3.33 | 0.32 | 1.00 |
| 105 | B marine - D terrestrial | 1.05 | 3.33 | 0.32 | 1.00 |
| 106 | C marine - D marine | 0.00 | 0.97 | 0.00 | 1.00 |
| 107 | C marine - A terrestrial | -4.58 | 3.31 | -1.38 | 0.99 |
| 108 | C marine - B terrestrial | -4.58 | 3.31 | -1.38 | 0.99 |
| 109 | C marine - C terrestrial | -3.51 | 3.31 | -1.06 | 1.00 |
| 110 | C marine - D terrestrial | -3.51 | 3.31 | -1.06 | 1.00 |
| 111 | D marine - A terrestrial | -4.58 | 3.32 | -1.38 | 0.99 |
| 112 | D marine - B terrestrial | -4.58 | 3.32 | -1.38 | 0.99 |
| 113 | D marine - C terrestrial | -3.51 | 3.32 | -1.06 | 1.00 |
| 114 | D marine - D terrestrial | -3.51 | 3.32 | -1.06 | 1.00 |
| 115 | A terrestrial - B terrestrial | 0.00 | 0.22 | 0.00 | 1.00 |
| 116 | A terrestrial - C terrestrial | 1.07 | 0.22 | 4.97 | **0.00** |
| 117 | A terrestrial - D terrestrial | 1.07 | 0.22 | 4.84 | **0.00** |
| 118 | B terrestrial - C terrestrial | 1.07 | 0.21 | 5.02 | **0.00** |
| 119 | B terrestrial - D terrestrial | 1.07 | 0.22 | 4.88 | **0.00** |
| 120 | C terrestrial - D terrestrial | 0.00 | 0.21 | 0.00 | 1.00 |

ST8. Parameter estimates from a Gaussian generalized linear mixed-effects model (log link) evaluating the effects of ecosystem type on future trait velocity. Fixed-effect estimates are shown for the conditional (mean) model, with freshwater ecosystem as reference level. The dispersion model reports effects of ecosystem on residual variance. Random effects represent among-study and among-taxon variation in intercepts (n = 821, study = 39, taxon = 70). Estimates are reported with standard errors, Wald z-values, and associated p-values. Significant p-values are in bold.

| **Effect** | **Component** | **Term** | **Estimate** | **Standard Error** | **Z-value** | **P-value** |
| --- | --- | --- | --- | --- | --- | --- |
| *Conditional Model* | |  |  |  |  |  |
| fixed | cond | (Intercept) | 3.628 | 0.909 | 3.992 | 0.000 |
| fixed | cond | ecosystemintertidal | 0.073 | 1.078 | 0.068 | 0.946 |
| fixed | cond | ecosystemmarine | 0.059 | 1.078 | 0.055 | 0.956 |
| fixed | cond | ecosystemterrestrial | 0.212 | 0.964 | 0.220 | 0.826 |
| *Dispersal Model* | |  |  |  |  |  |
| dispersion | | (Intercept) | -0.235 | 0.424 | -0.555 | 0.579 |
| dispersion | | ecosystemintertidal | 0.323 | 0.432 | 0.748 | 0.454 |
| dispersion | | ecosystemmarine | -0.208 | 0.443 | -0.470 | 0.638 |
| dispersion | | ecosystemterrestrial | 0.076 | 0.428 | 0.177 | 0.859 |
| *Conditional Model* | |  | **Variance** | **Standard Deviation** | |  |
| ran_pars | cond | sd__(Intercept) | 1.539 | 1.241 |  |  |
| ran_pars | cond | sd__(Intercept) | 0.289 | 0.537 |  |  |

ST9 Number of pairwise population comparisons (between populations with difference in UTL) between each realm that have less phenotypic cline than the projected increase in temperature at the site of the lowest population (excluded represents number of populations that have less). Included are the number of population pair-wise comparisons that have a difference in UTL between populations greater than the increase in warming at the population sites. Excluded is the number of comparisons that have less divergence relative to increase in habitat temperature.

| **Ecosystem** | **Included** | **Excluded** | **Total** |
| --- | --- | --- | --- |
| Freshwater | 1 | 9 | 10 |
| Intertidal | 148 | 87 | 235 |
| Marine | 44 | 47 | 91 |
| Terrestrial | 137 | 348 | 485 |
